# PICKLE recruits RETINOBLASTOMA RELATED 1 to Control Lateral Root Formation in *Arabidopsis*

**DOI:** 10.1101/643122

**Authors:** Krisztina Ötvös, Pál Miskolczi, Peter Marhavý, Alfredo Cruz-Ramírez, Eva Benková, Stéphanie Robert, László Bakó

## Abstract

Lateral root (LR) formation is an example of plant post-embryonic organogenesis event. LRs are issued from non-dividing cells entering consecutive steps of formative divisions, proliferation and elongation. The chromatin remodeling protein PICKLE negatively regulates auxin-mediated LR formation through a mechanism that is not yet known. Here we show that PICKLE interacts with RETINOBLASTOMA-RELATED 1 (RBR1) to repress the *LATERAL ORGAN BOUNDARIES-DOMAIN 16* (*LBD16*) promoter activity. Since LBD16 function is required for the formative division of LR founder cells, repression mediated by the PKL-RBR1 complex negatively regulates formative division and LR formation. Inhibition of LR formation by PKL-RBR1 is counteracted by auxin indicating that in addition to auxin-mediated transcriptional responses, the fine-tuned process of LR formation is also controlled at the chromatin level in an auxin-signaling dependent manner.

## Introduction

Lateral roots (LRs) are initiated in non-dividing pericycle cells by the plant hormone auxin that first triggers the reprogramming of non-dividing cells into proliferative LR founder cells, then later acts in a gradient for the execution of further steps in the LR developmental program. This developmental process initiates by auxin signaling converging on protoxylem pericycle cells, which promotes the degradation of AUXIN/INDOLE-3-ACETIC ACID (Aux/IAA) proteins involved in LR initiation (LRI) (Péret et al., 2012). Elimination of Aux/IAA repressors through SKP-Cullin-F-box (TIR1/AFB) (SCF^TIR1/AFB^) ubiquitin ligase complexes and the 26S proteasome results in activation of AUXIN RESPONSE FACTOR (ARF)7/ARF19 transcription factors to drive the expression of *LATERAL ORGAN BOUNDARIES DOMAIN/ASYMMETRIC LEAVES-LIKE (LBD/ASL) LBD16/ASL18*, *LBD29/ASL16* and many other target genes required for auxin response, LRI and development (Fukaki and Tasaka, 2008).

Shortly after auxin signal perception, a pair of pericycle cells adjacent to the xylem pole becomes polarized when nuclei of these cells migrate towards the common cell walls thereby creating intracellular asymmetry (De Smet and Beeckman, 2011). Anticlinal division of such polarized cells yields two larger flanking and two smaller central daughter cells, the latter of which continue to divide periclinally to form the LR primordia (Péret et al., 2012). Nuclear migration and establishment of asymmetry in LR founder cells is compromised in plants expressing a dominant negative version of LBD16 suggesting that LBD16 is one of the key players mediating formative cell division and LRI (Goh et al., 2012). Polar nuclear movement and anticlinal cell division is inhibited in the gain-of-function *solitary-root* (*slr-1)* mutant expressing a non-degradable version of the SLR/IAA14 repressor protein hence the mutant lacks lateral roots (De Rybel et al., 2010, Fukaki et al., 2002). Overexpression of CYCLIN D3;1, a known activating subunit of the G1/S regulator CDKA;1 kinase triggers a few rounds of pericycle division but fails to initiate LR formation in the *slr-1* root (Vanneste et al., 2005). Conversely, disruption of the *PICKLE* (*PKL*) gene encoding a Chromodomain-Helicase-DNA binding (CHD) ATP-dependent chromatin remodeling factor restores LR formation in the *slr-1* background indicating that inactivation of the *PKL* gene enables both the initial formative divisions as well as the subsequent organized proliferation of pericycle cells (Fukaki et al., 2006). It has been therefore proposed that PKL negatively regulates LR initiation at the chromatin level, however the mechanism through which PKL acts remained obscured.

PICKLE is a plant homologue of the animal chromatin remodeling ATPase Mi-2/CHD3/4 proteins which in vertebrates form the Mi-2/Nucleosome Remodeling and Deacetylase (NuRD) repressor complexes regulating chromatin organization, gene transcription and developmental signaling (Denslow and Wade 2007). Animal NuRD complexes contain the ATPase chromatin remodeler CHD3/CHD4 proteins and a histone deacetylase subcomplex that comprises the histone deacetylase HDAC1/HDAC2 enzymes and the retinoblastoma-binding RbAp46 and RbAp48 histone chaperon proteins (Allen et al., 2013). The presence of class 1-type histone deacetylases and a panel of RbAp46/48 homologues in the *Arabidopsis* genome suggests that similar to animal systems plant Mi-2/CHD3/4 ATPase remodelers might assemble to NuRD-like complexes. However, biochemical characterization of the *Arabidopsis* PKL protein failed to find evidence for the existence of such complexes thus far (Ho et al., 2012).

Intriguingly, the PKL protein sequence contains two LxCxE peptide motifs that is often present in viral and cellular proteins and mediates stable binding by fitting into a groove within the conserved small pocket domain of retinoblastoma (pRB) proteins (Lee et al., 1998, Singh et al., 2005). Animal retinoblastoma proteins and the plant ortholog RETINOBLASTOMA-RELATED 1 (RBR1) control the G1-to-S-phase progression in the cell cycle (Gutzat et al., 2012). In G1 phase the hypophosphorylated form of pRB binds to and inactivates the E2F/DP1 transcription factor heterodimer, the activity of which is necessary for G1-to-S progression. Phosphorylation of pRB by cyclin-dependent kinase (CDK)/CyclinD complexes releases active E2F/DP1 dimers, initiates the transcription of S-phase specific genes and triggers cell division. Several lines of evidence indicate that the function of retinoblastoma proteins extends much beyond the canonical G1-to-S-phase control role. Human pRB protein has been implicated in cellular differentiation by associating with tissue-specific transcription factors and modulating their activity (Poznic, 2009). In vertebrates pRB is often present in chromatin repressor complexes that are having roles in developmental transitions (Harbour and Dean, 2000). These findings strongly suggest that the pRB protein regulates cellular differentiation separate from its function in cell cycle progression (Berman et al., 2008). Plant RBR proteins share the basic structural and functional features of pRB (Gutzat et al., 2012, Desvoyes et al., 2014). Similar to animal pRB, plant RBR proteins can associate with histone deacetylases to repress gene transcription (Rossi et al., 2003). While human pRB binds to histone deacetylases directly through the LxCxE (Nicolas et al., 2000), plant HDAC proteins do not contain the LxCxE motif and accordingly, RBR proteins interact with HDACs indirectly. It has been reported that the *Arabidopsis* RBR1 binds to the MULTICOPY SUPPRESSOR OF IRA1 (MSI1) protein, which is a plant homologue of the animal RbAp46/48 proteins (Julien et al., 2008). Evidence indicates that members of the plant MSI protein family associate with histone deacetylases to mediate transcriptional silencing at target loci (Gu et al., 2011). The RBR1-MSI1 interaction takes place at the RbA pocket domain of RBR1 leaving the LxCxE binding cleft that is located on the RbB pocket domain available for protein binding (Julien et al., 2008). This interaction topology enables RBR1 to recruit histone deacetylases and simultaneously associate with transcription factors and chromatin modifiers containing the LxCxE motif.

We report here that PKL interacts with the RETINOBLASTOMA-RELATED 1 (RBR1) protein in the *Arabidopsis* root. Consistent with this finding we show that similar to the PKL protein, RBR1 is a negative regulator of LR formation. Our data further demonstrate that PKL recruits RBR1 to the promoter of the LR-specific *LBD16* gene. When bound to the promoter the PKL-RBR1 complex acts as a transcriptional repressor of *LBD16* and negatively regulates LR formation. Through the IAA14/ARF7/ARF19 signaling pathway auxin releases the PKL-RBR1 complex from the *LBD16* promoter indicating that this novel, chromatin level regulation of LR formation is tightly coupled to auxin signaling.

## Results

### PKL interacts with *Arabidopsis* RBR1

To test whether PKL and RBR1 proteins interact, we expressed epitope-tagged versions of PKL and RBR1 proteins in *Arabidopsis* protoplasts and analyzed their interaction by co-immunoprecipitation (co-IP) assays. Our results demonstrated that RBR1 binds to the PKL protein (Figure 1A). To confirm interaction by an independent method, we performed bimolecular fluorescence complementation (BiFC) assays. Interaction between RBR1 and the full-length PKL were detected in the nucleus (Figure 1B). Finally, the PKL/RBR1 interaction between the endogenous PKL and RBR1 proteins was validated *in planta* by co-IP experiment in five-day-old wild type seedling roots (Figure 1C). Since PKL harbours two LxCxE motifs we tested epitope-tagged, truncated versions of PKL for RBR1 interaction to find similar affinity binding of the N- and C-terminal halves (Figure S1A-C). We also examined whether mutation of the peptide motifs disrupts binding by converting both LxCxE motifs in the PKL sequence to AxAxA. BiFC interaction assays showed that mutations weakened but did not abolished interaction indicating that in addition to the LxCxE motif other domains of PKL are also involved in contacting the RBR1 protein (Figure S1D).

**Figure 1.**
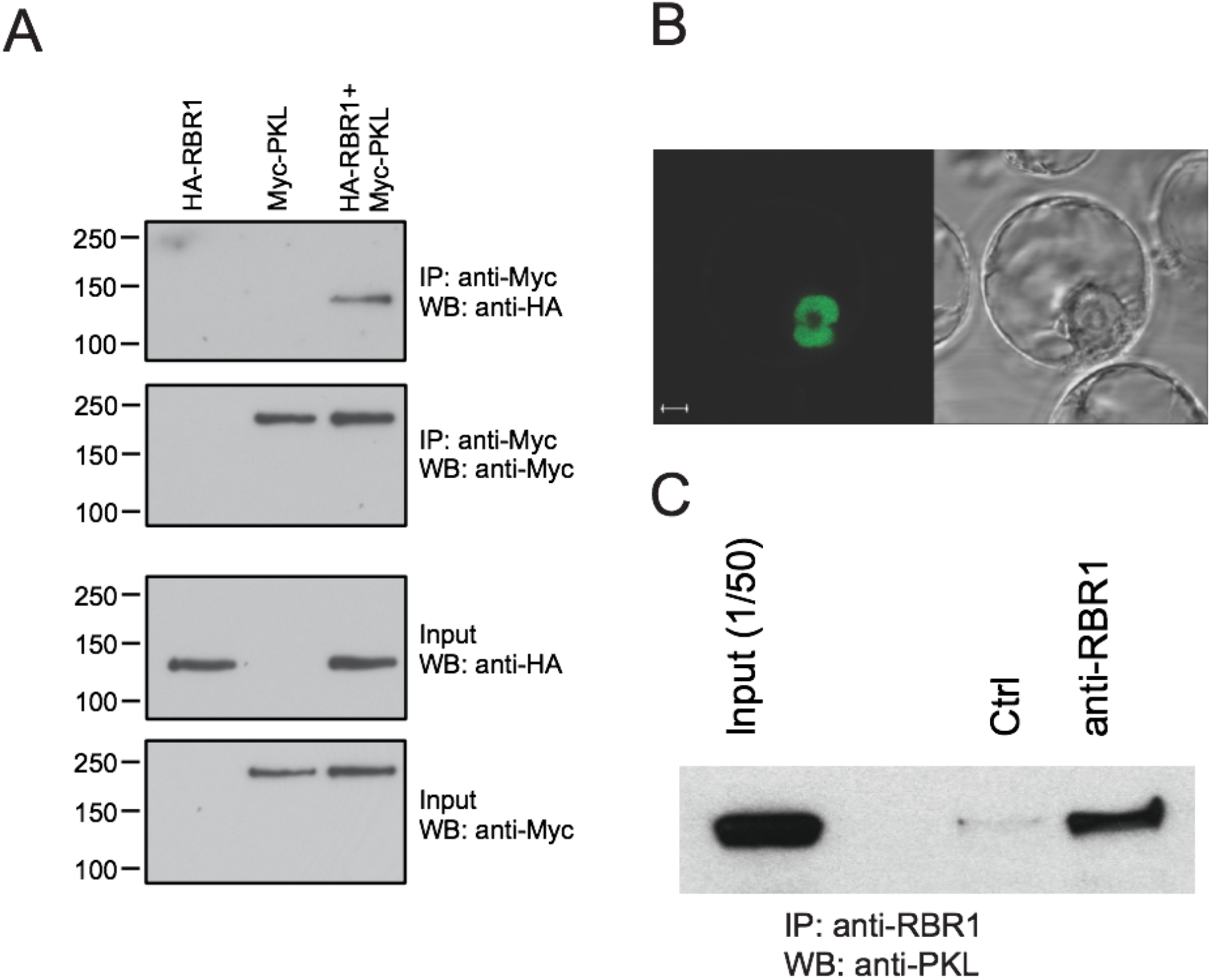
PKL interacts with RBR1 *in vivo*. (A) Protoplasts were transfected with *35S::HA-RBR1* (HA-RBR1) and *35S::myc-PKL* (Myc-PKL) constructs and protein extracts from transformed protoplasts were immunoprecipitated with anti-myc antibodies. Immunocomplexes and input proteins were analyzed on protein gel blots using anti-HA or anti-myc antibodies, respectively. (B) Confocal microscopic image of the subcellular localization of the RBR1/PKL complex by BiFC assays. Coexpression of *35S::YFPN-RBR1* (RBR1) and *35S::YFPC-fPKL* (fPKL) in suspension derived *Arabidopsis* protoplasts. Bar = 5 μm. (C) CoIP assay in 5 DAG wild type Col-0 roots. Protein extracts were immunoprecipitated either with preimmune serum (Ctrl) or with anti-RBR1 antibody and immunocomplexes were analyzed on protein gel blots using anti-PKL antibody. Input is 1/50th of the total protein amount used for the immunoprecipitation reactions.

### RBR1 is expressed in xylem pole pericycle cells

In *Arabidopsis* seedlings PKL is expressed in meristems, organ primordia and in the stele including the pericycle cells (Eshed et al., 1999; Fukaki et al., 2006, Figure S2) while RBR1 is abundant in proliferating tissues including the shoot and root apical meristems, proximal part of young leaves and emerging LRs (Borghi et al., 2010, Figure S2). To explore whether the RBR1 protein is present in pericycle cells where PKL presumably functions to repress LR formation, we introduced the *pRBR:RBR-RFP* construct into the enhancer trap line J0121 in which GFP expression is restricted to XPP cells (Laplaze et al., 2005). Confocal microscopic analysis of roots expressing the RBR1-RFP fusion protein revealed that RBR1 is present in xylem pole pericycle cells (Figure 2). Interaction of PKL with RBR1 in the *Arabidopsis* root and the overlapping expression pattern of the two proteins in the differentiation zone indicated that RBR1 might participate in the PKL mediated repression of LR formation.

**Figure 2.**
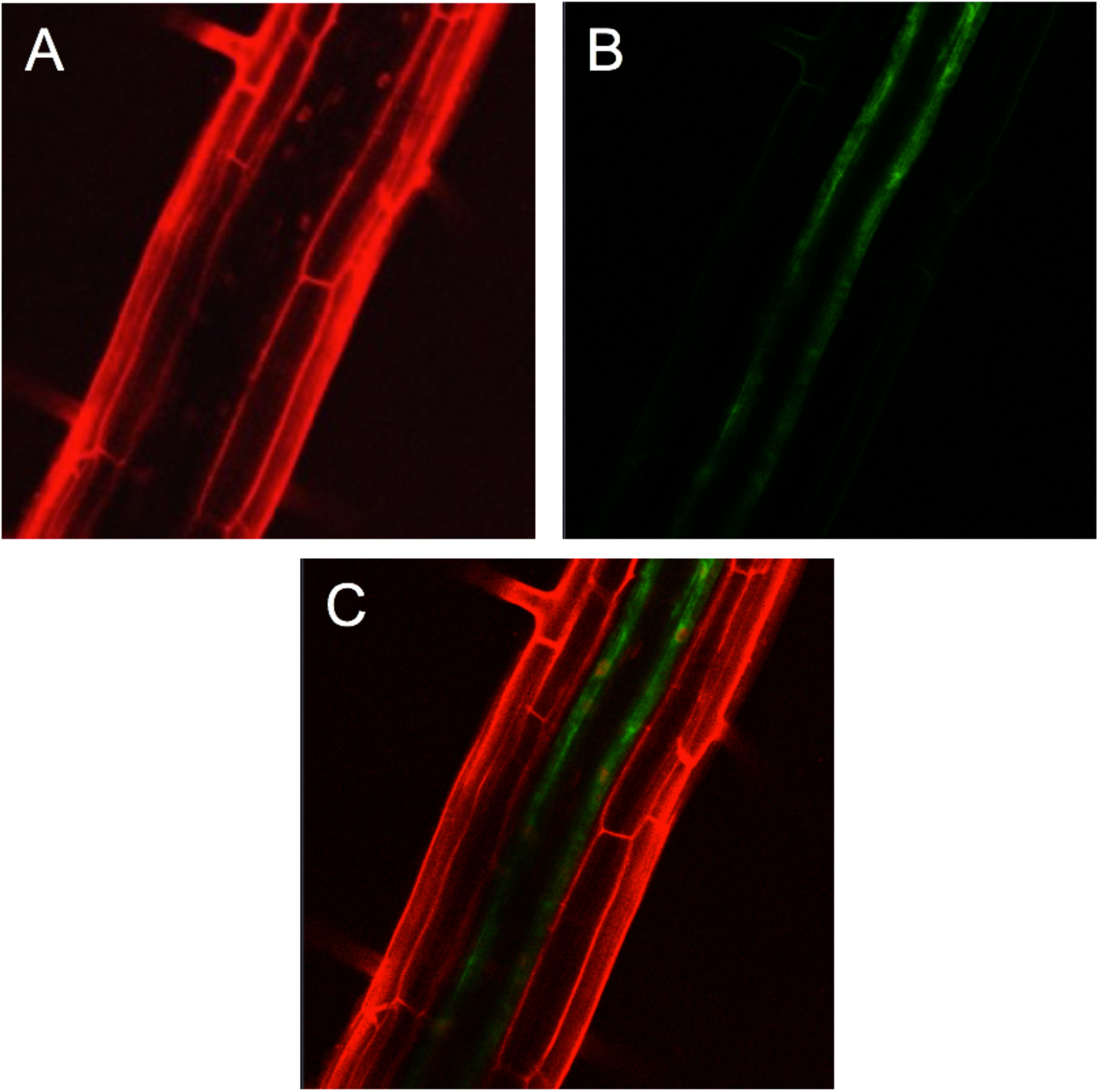
The RBR1 protein is present in xylem pole pericycle cells. Longitudinal confocal section of J0121::*pRBR:RBR-RFP* root showing (A) RBR-RFP fluorescence (red) counterstained with propidium iodide, (B) GFP fluorescence (green) marking XPP cells and (C) overlay of the two image.

### Similar to PKL, the RBR1 protein is a negative regulator LR formation

Previous characterization of the *pkl/ssl2-1* mutant expressing a short N-terminal fragment of the PKL protein showed that the *ssl2-1* root is significantly shorter but produces LRs at similar density than the wild type (Fukaki et al., 2006). However, under our experimental conditions phenotypic analysis of *ssl2-1* confirmed the shorter root phenotype but indicated a significantly higher LR density compared to the wild type (Figure 3A-B, Figure S3A, B). To assess if RBR1 had a role in LR development, we induced a reduction of RBR1 level in the diploid plant by partially silencing *RBR1* expression with the production of an artificial microRNA directed against the 3’ UTR of the *RBR1* mRNA (Cruz-Ramírez et al., 2013). Molecular analysis of the *amiRBR1* line showed that silencing decreased both transcript and protein levels by 50% compared to the wild type (Figure 3C-D). Phenotypic examination of the seedling roots revealed unperturbed primary root growth however an increased root branching was observed in the *amiRBR1* line (Figure 3 E-G). To test whether root branching was sensitive to RBR1 abundance, we increased RBR1 level by expressing the RBR1-RED FLUORESCENT PROTEIN (*pRBR:RBR-RFP*) (Ingouff et al., 2006). The presence of this construct in the *rbr2-1* background lead to an increased expression of the wild type gene possibly due to the positive autoregulatory function of RBR1 over its own expression (Park et al., 1994) (Figure 3C-D). Primary root growth was unaffected in the *RBR1-RFP* line while root branching decreased compared to the wild type seedling root (Figure 3 E-G). Collectively, our data support a negative role for RBR1 in LR formation and indicate that an inverse relationship exists between RBR1 expression level and LR density. The fact that lower levels of both RBR1 and/or PKL enhanced LR formation and the overlapped expression pattern of PKL and RBR1 (Figure S2) suggested a functional relationship between the two proteins in LR formation.

**Figure 3.**
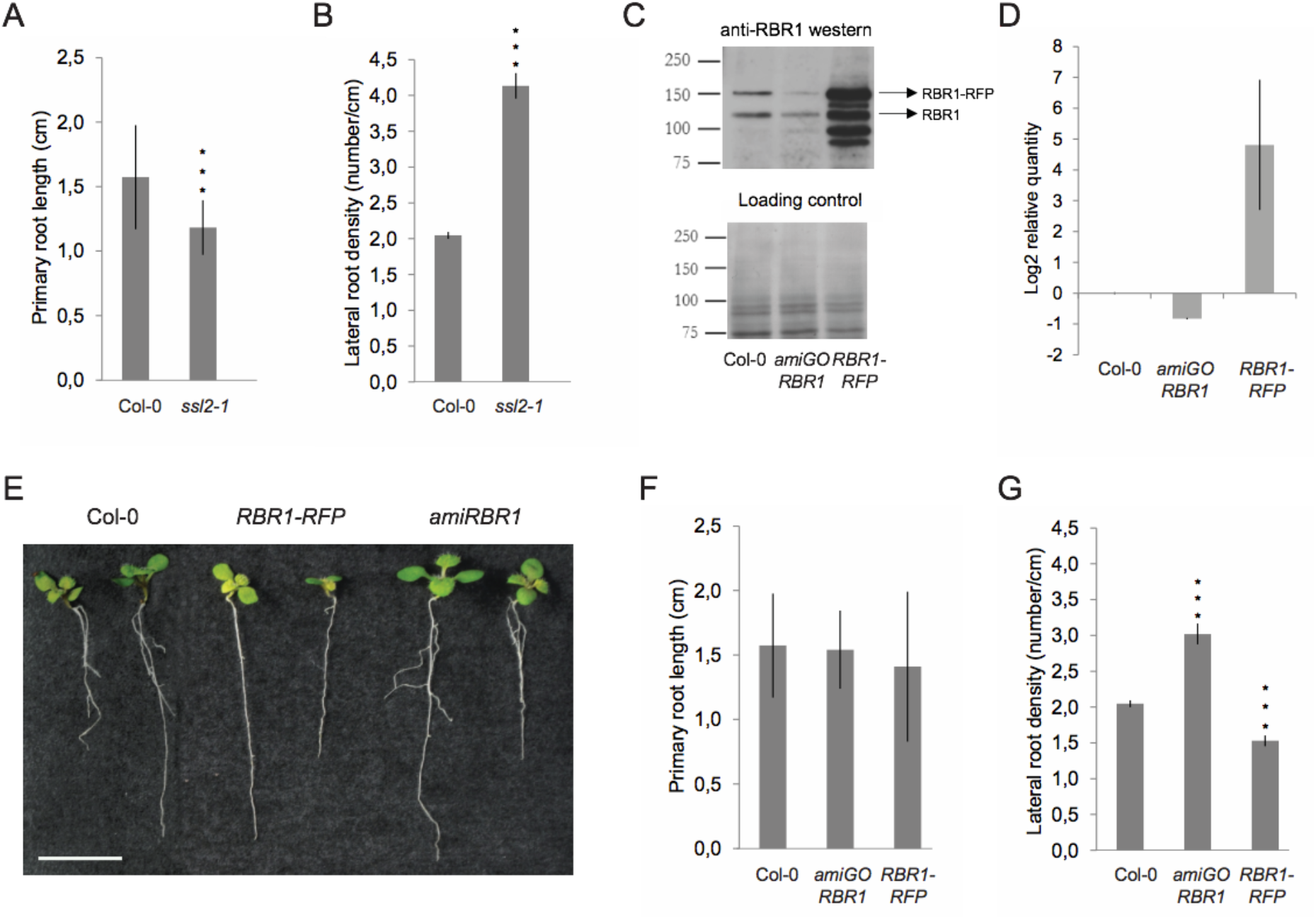
RBR1 protein abundance affects lateral root development. (A, B) Phenotypic analysis of 10 DAG *ssl2-1* roots. Quantification of the primary root length (A) and the lateral root density (B) in 10 DAG Col-0 and *ssl2-1* line. Error bars represent means ± SD of the mean. The data was normalized to the levels in Col-0 p < 0,001 by two-sided t-test; n_Col-0_ = 57, n_*ssl2-1*_ = 41. (C) Western blot analysis of RBR1 protein level and (D) quantitative real time RT-PCR (qRT-PCR) analysis of *RBR1* mRNA expression in wild type Columbia (Col-0), *amiRBR1* and *pRBR1::RBR1-RFP* (*RBR1-RFP*) lines. The expression level of *RBR1* was normalized to At1g61670 and At3g03210 (mean stability value M = 0.0405) and data (log2) from three technical replicates are shown as means ± standard deviation (SD). (E) Phenotype of 10 DAG seedlings expressing different levels of RBR1. Bar = 1 cm. (F) Quantification of the primary root length (cm) and (G) the lateral root density in 10 DAG Col-0, *amiRBR1* and *RBR1-RFP* roots. Error bars represent means ± SD of the mean. The data was normalized to the levels in Col-0 p < 0,001 by two-sided t-test; n_Col-0_ = 57, n_*amiRBR1*_ = 50, n_*RBR1-RFP*_ = 52.

### PKL recruits RBR1 to the *LBD16* promoter

Polar nuclear movement and asymmetric divisions are blocked in LR founder cells of the *slr-1* mutant (De Rybel et al., 2010). Since disruption of *PKL* restored LR formation in the *slr-1* root, the PKL protein probably acts to repress either polar nuclear movement or the subsequent asymmetric cell divisions. Consistent with a PKL-RBR1 functional interaction, the *RBR1-RFP* line produced fewer LRs but did not show aborted LR initiation events indicating that RBR1 protein level also affects LRI at an early stage. Recent experimental data indicate that nuclear movement is mediated by the LBD16/ASL18 and related LBD/ASL transcription factors in the *Arabidopsis* root. Expression of a dominant negative repressor version of the LBD16 protein did not affect LR founder cell specification but prevented nuclear migration and abolished LR formation (Goh et al., 2012).

To examine whether the PKL and RBR1 proteins are involved in the transcriptional regulation of the *LBD* genes that are known to have a role in LR initiation, we performed chromatin immunoprecipitation (ChIP) assays. We designed primers specific for the promoter of *LBD16*, *LBD17*, *LBD18*, *LBD29* and *LBD33* genes and analyzed RBR1-bound chromatin samples by PCR. Out of the five *LBD* genes tested, ChIP with the RBR1 antibody pulled down DNA fragments of the *LBD16* promoter only (Figure 4A and Figure S4). Analysis of the *LBD16* promoter sequence revealed the presence of a consensus TTTGCCGG E2F binding motif 1374 bp upstream of the translation initiation site. Similar to pRB proteins of human and fly, *Arabidopsis* RBR1 binds to members of the E2F transcription factor family (Magyar et al., 2012) and can be recruited to promoters containing the consensus E2F binding site sequence (Weimer et al., 2012; Zhao et al., 2012). We therefore tested if the interaction of RBR1 with the *LBD16* promoter was mediated by E2F transcription factor. Our data showed this was not the case, fragment containing the predicted E2F binding site could not be detected in ChIP samples. By contrast, the promoter proximal fragments could be amplified indicating that binding takes place close to the transcription start site (TSS) (Fragments F2 and F4 in Figure 4A).

**Figure 4.**
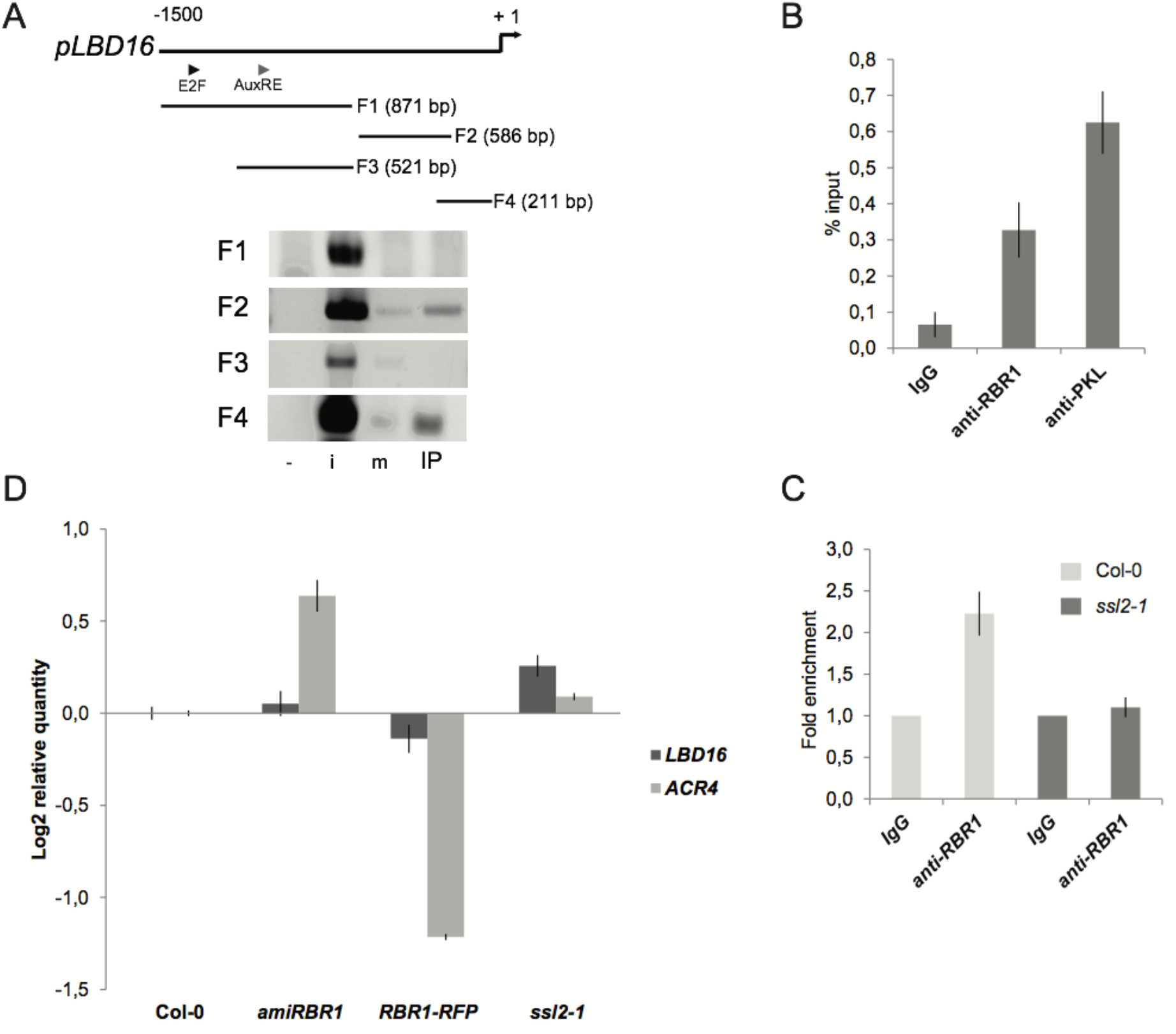
PKL recruits RBR1 to the *LBD16* promoter. (A) ChIP-PCR analysis of different *LBD16* promoter fragments (F1-F4) precipitated with RBR1 antibody using chromatin extracted from 10 DAG Col-0 roots. Triangles show the position of the predicted E2F consensus (E2F) binding site, the AuxRE motif and the transcription start site (TSS), +1 labels the translation initiation site. -: non-template control, i: input DNA, m: IP with IgG, IP: IP with anti-RBR1 antibody. (B) ChIP-qRT-PCR analysis of RBR1 and PKL binding to *LBD16* promoter fragment (F4) in 5 DAG wild type Columbia roots. Bar graphs show quantification of qRT-PCR products from ChIP experiment using anti-RBR1 and anti-PKL antibody, expressed as % input. Data from two biological replicates were shown as means ± SD of the mean. (C) ChIP-qRT-PCR analysis of RBR1 binding to the *LBD16* promoter in *ssl2-1* background. Bar graphs show quantification of qRT-PCR products from ChIP experiment using anti-RBR1 antibody on chromatin extracted from 10 DAG Col-0 and *ssl2-1* roots. Data from two biological replicates were shown as means ± SD of the mean. (D) qRT-PCR analysis of *LBD16* and *ACR4* gene expression in 10 DAG roots expressing *35S::amiRBR1* (*amiRBR1*), *pRBR1::RBR1-RFP* (*RBR1-RFP*) and in the *PKL* mutant *ssl2-1* compared to Col-0. The expression levels was normalized to At1g61670 and At3g03210 (mean stability value M = 0.0413) and data (log2) from three technical replicates were shown as means ± SD.

Interaction between PKL and RBR1 proteins suggested they act on a common pathway to regulate LR formation, thus the presence of PKL on the *LBD16* promoter has been tested by ChIP assay. Quantitative real-time PCR (qRT-PCR) analysis of PKL-bound chromatin samples revealed that PKL binds to the *LBD16* promoter and binding takes place within the same proximal promoter region (Fragment F4) to which RBR1 binds (Figure 4B). To assess whether RBR1 binds directly or through interaction with PKL to the *LBD16* promoter we examined promoter binding in the *ssl2-1* root that lacks the full-length PKL protein (Figure S3B). ChIP with the anti-RBR1 antibody failed to show any enrichment of the *LBD16* promoter fragment (Figure 4C). That PKL interacts with RBR1 and promoter targeting is abolished when the PKL protein is absent collectively indicate that binding of the complex to the *LBD16* promoter takes place through PKL.

### Transcriptional and functional analysis of the effect of RBR1 and PKL proteins on *LBD16* expression

To assess the effect of PKL and RBR1 proteins on the *LBD16* promoter activity we quantified *LBD16* gene expression in wild type and mutant roots. *LBD16* expression was increased in the *amiRBR1* and *ssl2-1* lines and attenuated in the *RBR1-RFP* line, suggesting that RBR1 and PKL proteins act as repressors of the *LBD16* promoter activity (Figure 4D). The observed negative correlation between the abundance of PKL and RBR1 proteins and *LBD16* gene expression led us to test whether a similar relationship exists between protein levels and asymmetric cell division. Asymmetric divisions in roots of *amiRBR1*, *RBR1-RFP* and *ssl2-1* lines were quantified by analyzing the expression of the *ARABIDOPSIS CRINKLY4* (*ACR4)* gene that is a marker of formative pericycle divisions during LRI (De Smet et al., 2008). *ACR4* transcript levels were increased in the *amiRBR1* and *ssl2-1* lines while decreased in the *RBR1-RFP* line (Figure 4D) indicating that by binding to the *LBD16* promoter PKL and RBR1 proteins ultimately regulate asymmetric cell division at LRI.

To exclude the possibility that altered RBR1 and PKL abundance would result in misexpression of *LBD16*, we introduced the *pLBD16:GFP* construct into the *amiRBR1*, *RBR1-RFP* and *ssl2-1* lines. The *LBD16* expression domain was unaffected (Figure S5) however the basal activity of the *LBD16* promoter in the *amiRBR1* and *ssl2-1* lines was strongly increased along the stele (Figure 5A). Consistent with the qRT-PCR (Figure 4D) and the LR density data (Figure 1B and 3G), elevated *LBD16* promoter activity gave rise to increased LR density (Figure 5A, asterisks). Higher basal activity of the *LBD16* promoter in the *amiRBR1* and *ssl2-1* lines might be due to an enhanced auxin activation response brought about by a less repressed state of the promoter. To test this hypothesis, we performed root bending assays to change the auxin distribution and concentration which in turn induces LR formation as a consequence of the gravitropic stimulus (Lucas et al., 2008; De Smet et al., 2007). Expression of the GFP reporter was much stronger in *amiRBR1* and *ssl2-1* background compared to wild type indicating that activation of the *LBD16* promoter is enhanced when either RBR1 or PKL protein levels are low and the promoter is not subjected to repression at the chromatin level (Figure 5B and C).

**Figure 5.**
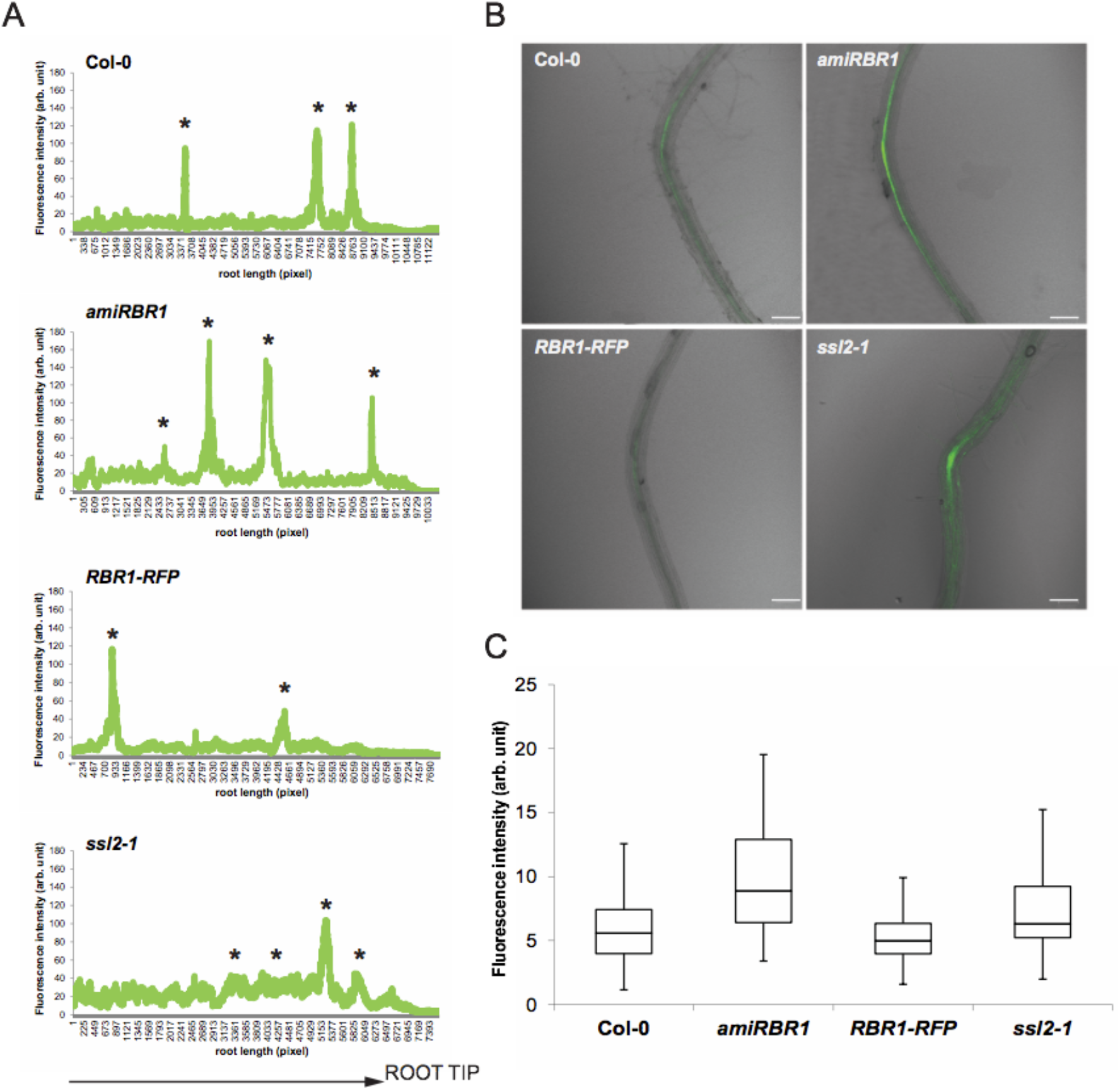
Functional analysis of the effect of RBR1 and PKL proteins on *LBD16* promoter. (A) Activity of the *pLBD16:GFP* reporter construct in 6 DAG roots of Col-0, *amiRBR1*, *RBR1-RFP* and *ssl2-1* lines. The graph shows GFP fluorescence intensity (arbitrary units) along the stele versus root length (in pixels). Asterisk (*) represents lateral root primordia/lateral roots. (B) Result of the root bending test. The confocal images show representative pictures from the bended sites of roots of different lines expressing *pLBD16::GFP* after 5 h following the 12-h-long gravistimuli and (C) the quantified data are summarized in box plots. The intensity of GFP fluorescence (arbitrary units) was measured in at least 31 roots per line. n_Col-0_=39, n_*amiRBR1*_=41, n_*RBR1-RFP*_=31, n_*ssl2-1*_=35.

### Chromatin context is required for proper control of *LBD16* promoter activity

To study PKL-RBR1 repressor binding to the *LBD16* promoter in a system where auxin levels can be easily manipulated, we expressed a *pLBD16:GFP* reporter construct in suspension culture-derived protoplasts. In the absence of auxin, a strong basal activity of the *LBD16* promoter was detected and this basal activity was not induced when transfected cells were cultured in the presence of auxin (Figure 6A). By contrast, the well-established auxin-signaling reporter *pDR5rev:GFP* (Ulmasov et al., 1997) showed the expected auxin inducible expression pattern in transfected protoplasts (Figure 6A). Activity of the *LBD29* promoter, another LR specific gene, as well as expression of the endogenous *LBD16* gene were also inducible by auxin indicating that the auxin signaling pathway is functional in transfected protoplasts (Figures 6A and D). Deletion of the distal part of the *LBD16* promoter (from −1547 to −811bp) containing binding sites for E2F and ARF transcription factors decreased its activity by about 60% while deletion of the proximal part of the promoter (from −811 to +1bp) reduced its activity by 90% relative to the full-length promoter (Figure 6B). Our prior data indicated that the PKL-RBR1 complex binds to the *LBD16* promoter and represses *LBD16* expression to control LR formation. Therefore, the high basal activity of the *pLBD16:GFP* transgene in transfected protoplasts could be due to the failure of the PKL-RBR1 repressor complex to bind the plasmid resident *LBD16* promoter. To test this hypothesis, we conducted transient plasmid ChIP assays (Lavrrar and Farnham, 2004) and analyzed the recovered DNA samples by using primers specific either to the plasmid-born or to the genomic *LBD16* promoter. Compared to the genomic promoter, almost no RBR1 binding was detectable on the extrachromosomal promoter (Figure 6C) indicating that the PKL-RBR1 repressor complex does not bind to the plasmid-born promoter. Overall these data demonstrate that outside of its chromatin context *LBD16* promoter activation is independent of an auxin-mediated signaling pathway and suggest that the activity and auxin responsiveness of the *LBD16* promoter is regulated at the chromatin level in its genomic context.

**Figure 6.**
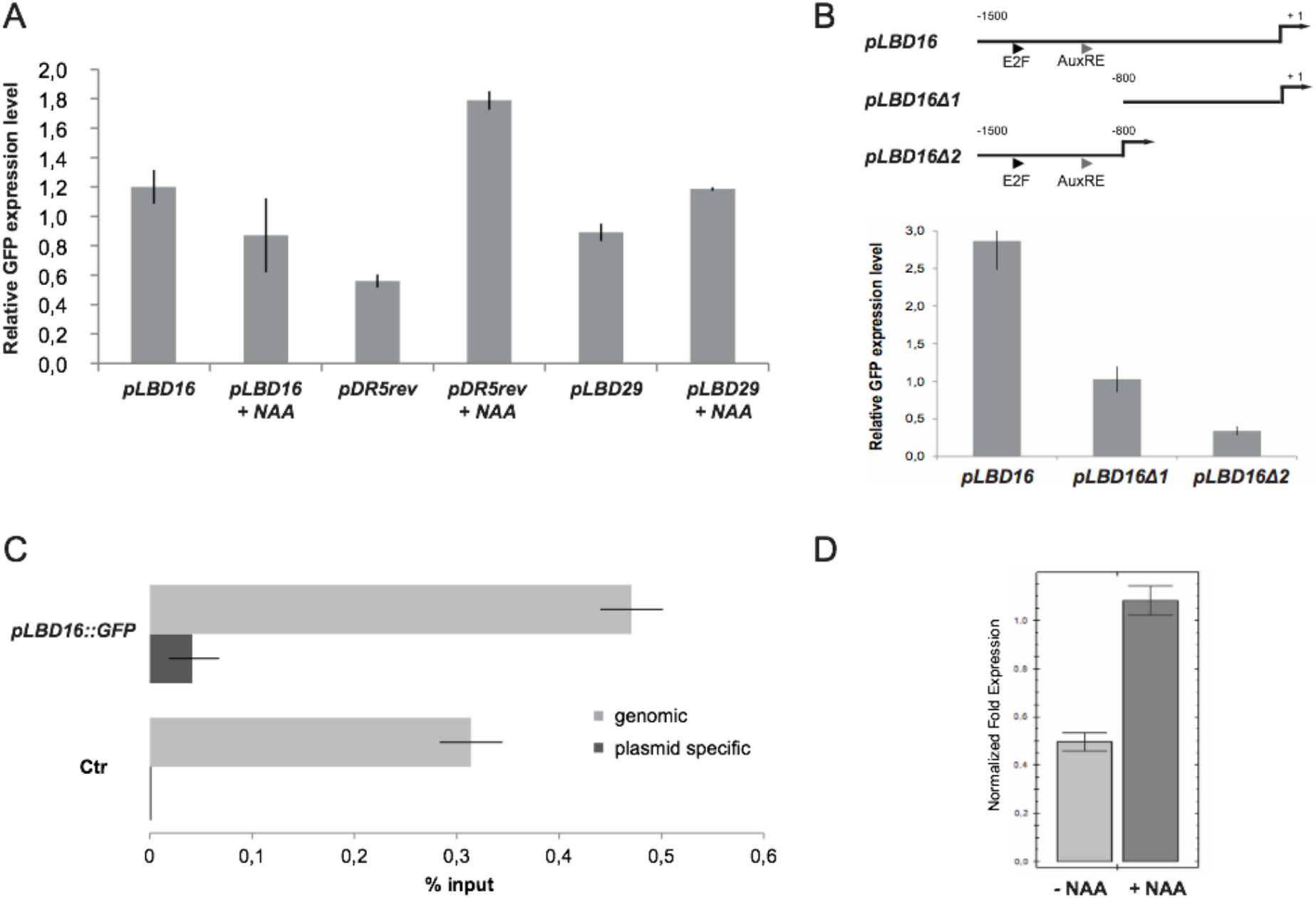
The *LBD16* promoter-reporter construct shows high and auxin-independent activity in *Arabidopsis* protoplasts. (A) GFP protein expression levels in suspension derived protoplasts treated with or without 10 μM NAA expressing the auxin responsive reporter *pDR5rev::GFP*, the lateral root specific *pLBD16::GFP* and *pLBD29::GFP* reporter constructs. Data from at least three biological replicates are shown as means ± SD of the mean. (B) ***LBD16*** promoter deletion assay. GFP protein expression levels in suspension derived protoplasts expressing the *pLBD16::GFP* (*pLBD16*), the *pLBD16Δ1::GFP* (*pLBD16Δ1*) and *pLBD16Δ2::GFP* (*pLBD16Δ2*) constructs. The deleted regions are shown in the diagram. Triangles mark the position of the predicted E2F consensus (E2F) and the AuxRE motives, +1 labels the translation initiation site. (C) ChIP-qRT-PCR analysis of RBR1 binding to genomic and plasmid-born *LBD16* promoter using chromatin isolated from protoplasts transfected either with control (Ctr) or with *pLBD16::GFP* construct. Bar graphs show quantification of qRT-PCR products from ChIP experiment using anti-RBR1 antibody, expressed as % input. Data from two biological replicates are shown as means ± SD of the mean. (D) qRT-PCR analyses of the endogenous *LBD16* gene expression level in the presence and in the absence of 10 μM NAA. The expression level of *LBD16* was normalized to At2g32760 and At3g03210 (mean stability value M = 0.0398) and data (linear) from three technical replicates were shown as means ± SD.

### Auxin signaling is required to dissociate the PKL-RBR1 complex from the *LBD16* promoter

Our experimental data indicated that PKL and RBR1 proteins interact in the *Arabidopsis* root and the complex associates with the *LBD16* promoter to repress *LBD16* gene expression and asymmetric cell division in the pericycle. According to the current model, *LBD16* promoter activity is regulated by auxin signaling through the SCF^TIR1^/IAA14/ARF7/19 pathway. How the PKL-RBR1 mediated repression mechanism integrates into this canonical model of transcriptional control and how auxin signaling is linked with chromatin level regulation of *LBD16* gene expression was unclear. To address these questions, we first tested whether TIR1/AFB receptor function is required by blocking the formation of the TIR1–IAA–Aux/IAA complexes with the auxin antagonist auxinol (Hayashi et al., 2012). Treatment of seedlings with 20 µM auxinol entirely prevented LRI in all lines indicating that even if limiting PKL or RBR1 protein levels precludes repressor complex formation, activation of the *LBD16* promoter and LRI still requires auxin action (Figure S7).

Next we investigated how the PKL-RBR1 mediated repression of the *LBD16* promoter responds to auxin signaling. We took advantage of the LR-inducible system in which application of exogenous auxin activates the LR initiation program in all pericycle cells (Himanen et al., 2002) and consequently, induces *LBD16* expression in a synchronized manner (Figure 7A). Activation of LRI enabled us to examine changes of *LBD16* promoter occupancy by using molecular tools. Treatment of 10 DAG wild type seedling roots with 10 µM of the synthetic auxin 1-Naphtaleneacetic acid (NAA) for 2 h reduced binding of RBR1 and PKL proteins to the *LBD16* promoter by 50% (Figure 7B). Conversely, we could only detect subtle changes in the dominant negative auxin signaling mutant *slr-1* root upon auxin treatment. Because of the lower expression level of RBR1 and PKL in the *slr-1* root (Figure S3D) binding to the *LBD16* promoter appeared generally weaker. Our data indicate that auxin signaling dissociates the RBR1-PKL complex from the *LBD16* promoter and that *LBD16* expression depends on the coordinated action of chromatin remodeling factors and auxin signaling pathway.

**Figure 7.**
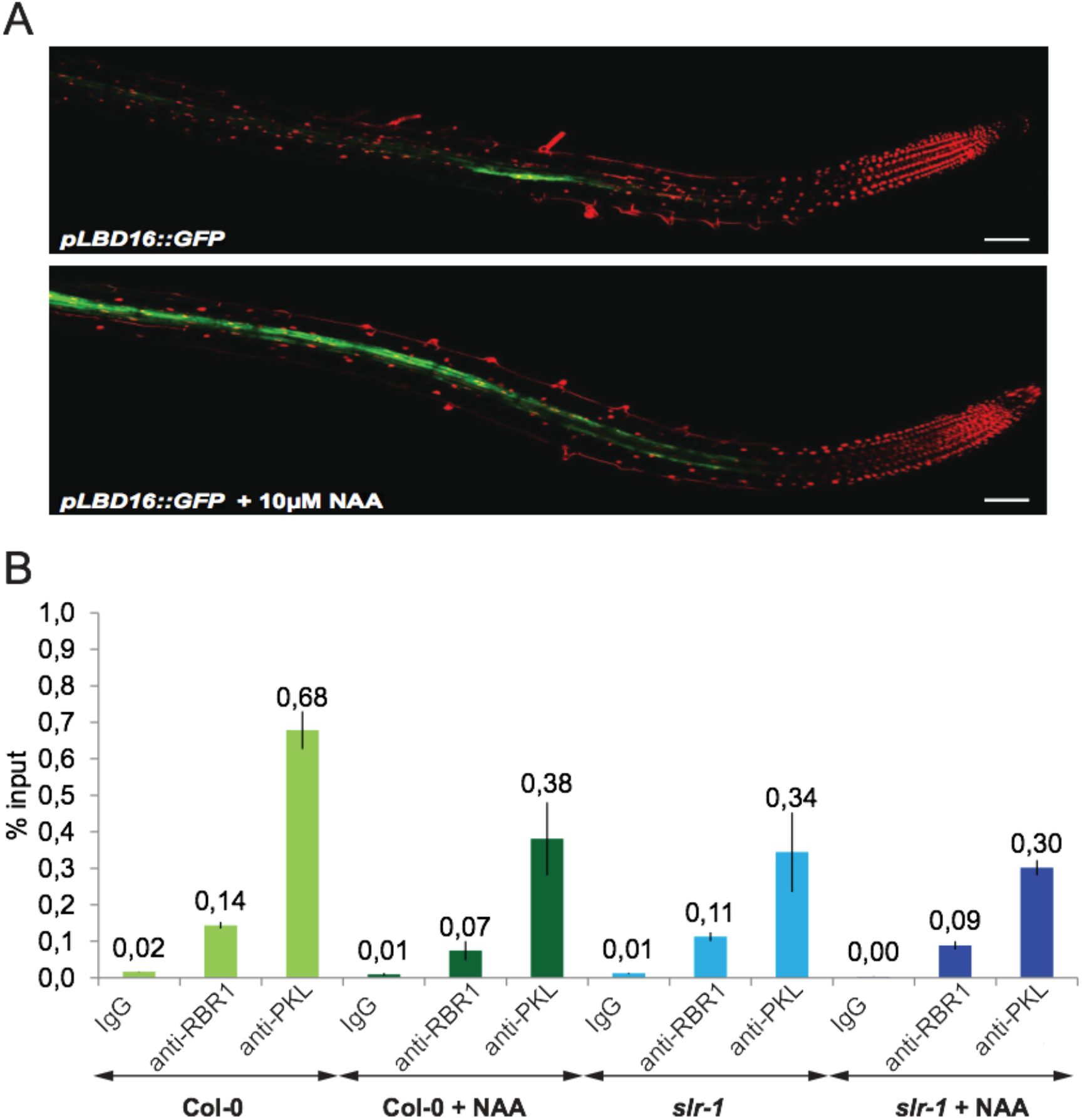
*LBD16* promoter activation upon NAA treatment. (A) Confocal microscopic images of 10 DAG Col-0 roots expressing the *pLBD16::GFP* construct with or without 10 μM NAA. (B) ChIP-qRT-PCR analysis of changes of RBR1 and PKL binding to *LBD16* promoter fragment (F4) in 10 DAG Col-0 and *slr-1* roots upon 10 μM NAA exposure. Bar graphs show quantification of qRT-PCR products from ChIP experiment using anti-RBR1 and anti-PKL antibodies, expressed as % input. Error bars represent means ± SD of the mean of three technical replicates.

## Discussion

Previous studies have identified the essential signaling response modules through which the phytohormone auxin initiates LR formation. The first module consists of the SLR/IAA14-ARF7-ARF19 proteins and reactivates the cell cycle as well as controls the first asymmetric divisions of the pericycle. The BODENLOS (BDL)/IAA12-MONOPTEROS (MP)/ARF5 pair forms the second auxin response module that shares some of the functions of the first module but also acts to prevent further unsolicited formative divisions (De Smet et al., 2010). In both modules binding of auxin to the TIR1/AFB receptors results in degradation of the Aux/IAA repressors of cognate ARF transcription factors and activates gene transcription. We show here that parallel to the destabilization of signaling repressors auxin also acts to disengage a repressor mechanism that controls *LBD16* promoter activity at the chromatin level. This repressor mechanism contains the ATP-dependent chromatin remodeler PKL that associates with the RBR1 protein in the root. The PKL-RBR1 complex binds to and represses the activity of the LR-specific *LBD16* gene promoter in an auxin-dependent manner and in turn restricts the first asymmetric cell divisions of the pericycle. Our data establish a link between auxin signaling and chromatin level regulation and reveals a novel mechanism underlying LRI in *Arabidopsis*.

### PKL recruits RBR1 to the *LBD16* promoter to form a repressive PKL-RBR1 complex

As an ATP-dependent chromatin remodeler of the CHD family the PKL protein sequence harbours PHD finger and chromo domains as well as SANT-SLIDE domains (Figure S1A) (Ho et al., 2013), each of which has been implicated in chromatin binding (de la Cruz et al., 2005). The N-terminus of the PKL protein contains two tandem chromo domains that have been proposed to bind methylated lysines of the histone H3 tail including H3K27me3 (Flanagan et al., 2005). In animals and plants the Polycomb repressive complex 2 (PRC2) mediates H3K27 trimethylation that is an important repressive mark with critical roles in developmental processes (Zheng and Chen, 2011). In *pkl* plants deposition of the H3K27me3 mark is significantly reduced suggesting that PKL itself somehow promotes H3K27me3 modification at target loci (Zhang et al., 2008). Recent ChIP data also show that PKL is present at these target loci however many of these genes show PKL independent expression (Zhang et al., 2012). Genome-wide mapping of histone H3K27me3 marks in *Arabidopsis* indicates that *LBD16*, *LBD17*, *LBD18*, *LBD29* and *LBD33* loci are all decorated by H3K27me3 (Roudier et al., 2011; Zhang et al., 2007). That in ChIP assays we detected binding of the PKL-RBR1 complex exclusively to the *LBD16* promoter suggests that targeting involves a mechanism other than the histone code reader function of the PKL chromo domain. Recently, PKL has been identified as a negative regulator of photomorphogenesis by binding to the transcription factors ELONGATED HYPOCOTYL5 (HY5)/HY5-HOMOLOG (HYH) (Jing et al., 2013). Targeting of PKL to promoters of cell elongation-related genes is compromised in the *hy5hyh* double knock-out mutant indicating that notwithstanding its DNA binding domains recruitment of PKL to definite gene loci requires association with sequence-specific transcription factors. We thus hypothesize that the observed selectivity of the RBR1-PKL complex for the *LBD16* promoter might also be mediated by a PKL-associated transcription factor. Our ChIP assays showed enrichment of F2 and F4 genomic fragments that are encompassing the −790 - +36 bp region from the proximal part of the promoter indicating that binding of the complex takes place near the TSS. Positioning of nucleosomes around the 5’ end of many eukaryotic genes shares common organization features according to which the TSS is located in a nucleosome-depleted region flanked by two arrays of nucleosomes (Yen et al., 2012). Pattern of nucleosome spacing and distribution is established and maintained by ATP-dependent chromatin remodeler complexes. Activity of the CHD1 chromatin remodelers Hrp1 and Hrp3 is required in the fission yeast to maintain nucleosome organization in gene coding regions (Hennig et al., 2012) whereas in the budding yeast the CHD3-CHD4 remodeler Mit1 plays a similar role (Lantermann et al., 2010). In *Arabidopsis* the SWI/SNF chromatin remodeling ATPase BRAHMA (BRM) regulates abscisic acid responses. In the absence of ABA signaling, BRM keeps the ABA-responsive *ABI3* and *ABI5* genes in a repressed state by binding to their promoter close to the TSS. Accordingly, BRM has been proposed to facilitate high occupancy of the +1 nucleosome adjacent to the TSS at the *ABI5* locus (Han et al., 2012). The TSS proximal occupancy of the PKL-RBR1 complex indicates that PKL might act in a similar manner to establish a repressive nucleosomal landscape at the *LBD16* locus. Positioning of the PKL-RBR1 complex is also consistent with recent genome-wide analyses of Rbf1 binding sites in *Drosophila* and RBR1 *in* Arabidopsis showing strong promoter-proximal targeting bias for both proteins (Acharya et al., 2012, Bouyer et al., 2018).

### Auxin signaling regulates *LBD16* expression by two distinct but interconnected mechanisms

Because LBD16 function mediates the first asymmetric divisions of pericycle cells, the mechanism controlling *LBD16* gene expression is of great importance for LR formation. Prior data have shown that auxin signaling initiates degradation of IAA14 repressor protein and in turn promotes dimerization of ARF7-ARF19 transcription factors that ultimately drive *LBD16* expression by binding to AuxRE cis-elements on the promoter (Okushima et al., 2007). This mechanism of promoter regulation can work as an on-off switch provided that the region between enhancer and TSS and in particular the TSS-adjacent core promoter is inherently silent in the absence of auxin. Our data however show that the proximal part of the *LBD16* promoter possesses fairly high basal activity indicating that the core promoter encompassing the TSS can initiate transcription without the trans-acting function of ARFs. Therefore, an additional, auxin responsive molecular mechanism must be present to keep tight control over the basal activity of the core promoter. We propose that the PKL-RBR1 complex fulfills this role. Both PKL and RBR1 proteins bind to the same proximal part of the promoter, the two proteins associate *in vivo* and the complex represses *LBD16* transcription and LR formation.

We further show that chromatin level regulation of the *LBD16* promoter activity acts in concert with auxin signaling as reduced PKL or RBR1 level alone is not sufficient for LR initiation, derepression of the *LBD16* promoter requires auxin and a functional SLR/ARF7/19 auxin signaling mechanism. Our data support that parallel to the SLR/ARF7/19 transcriptional module expression of the *LBD16* gene is controlled by the chromatin-bound PKL-RBR1 repressor complex. This dual regulation ensures that critical asymmetric cell divisions occur only upon auxin signal perception that is strictly coordinated with chromatin remodeling activities. The PKL-RBR1 complex represses *LBD16* promoter activity in an auxin-dependent fashion and in turn regulates asymmetric cell division. PKL binds either directly or through a sequence-specific transcription factor to the *LBD16* promoter. In the absence of auxin signaling PKL binds RBR1 that can recruit a chromatin modifier to the complex. Auxin signaling dissociates the complex probably by inducing post-translational modifications of either PKL or RBR1 proteins. RBR1 is hyperphosphorylated at the onset of LR formation (our unpublished data) and recent phosphoproteomic analysis of auxin-induced LR formation in *Arabidopsis* revealed that PKL is a phosphoprotein and its phosphorylation status changes upon application of the phytohormone (Zhang et al., 2013). It has been reported that the *Drosophila* dMi-2/CHD3 protein is phosphorylated by casein kinase 2 (dCK2) at the N-terminus and phosphorylation modulates its nucleosome binding and ATP-dependent nucleosome mobilization activities (Bouazoune and Brehm, 2005). Remarkably, the PKL N-terminal sequence contains a consensus CK2 phosphorylation site within the PHD domain and phosphoproteomic data indicates this site is phosphorylated in the root (Zhang et al., 2013). It is thus possible that similar to dMi-2, PKL activity and chromatin association is also under post-translational control. We observed that a fraction of PKL remains bound to the promoter, perhaps to prevent deposition of H3K27me3 marks or alternatively, due to its ATP-dependent remodeling activity PKL alters position of nucleosomes. Our findings presented here give the first insight on how chromatin level *regulation* of a key developmental gene is integrated with auxin signaling to control formative cell divisions and lateral organ formation

## Experimental Procedures

### Plant Material, Growth Conditions and Generation of Transgenic Lines

Seeds of *Arabidopsis thaliana* Columbia-0 wild type, *amiRBR1* (Cruz-Ramirez et al., 2012), *RBR1-RFP* (Ingouff et al., 2006), *ssl2-1* (Fukaki et al., 2006), *slr-1* (Fukaki et al., 2002), *pPKL:*:*PKL-GFP* in *pklpkr2* background (Aichinger et al., 2011) *and pLBD16::GFP* in Columbia-0, *amiRBR1*, *RBR1-RFP* and *ssl2-1* background were used. All seeds were grown on soil at 22°C 16 h light (150 µmol m^−2^ s^−1^) and 18°C (8 h dark) at 60% relative humidity. For aseptic growth, seeds were sterilized for 1 min with 70% (v/v) ethanol, soaked in NaOCl solution (1.2 % NaOCl, 0.05% Triton X-100) for 7 min, washed three times with sterile deionized water, and plated on 1% agar plates containing 0.5X Murashige and Skoog salt mixture including vitamins (Duchefa) and 0.5% Sucrose. The plates were maintained in darkness at 4°C for 2 days for stratification and then placed for 10 days at 22°C 16 h light (150 µmol m^−2^ s^−1^) and 18°C (8 h dark) at 60% relative humidity. 5 DAG, roots in vast quantity were harvested by cutting the roots of 5 DAG seedlings produced by using a hydroponic culture system, kept at 22°C 16 h light (150 µmol m^−2^ s^−1^) and 18°C (8 h dark) at 60% relative humidity. Transgenic *Arabidopsis* plants were generated by *Agrobacterium* mediated transformation using the floral dip method (Clough and Bent, 1998).

### Generation of plasmid constructs

All plasmid constructs were made using standard cloning techniques and polymerase chain reaction (PCR). Original cDNA clones were purchased from the Riken cDNA collection (http://www.brc.riken.jp). The N-terminal part of the PKL protein (aa 1-586) was synthetized by GenScript (http://www.genscript.com). Details of the molecular cloning work are provided in the Supplementary Experimental Procedures.

### Production of antibodies

Anti-RBR1 antibody has been generated using a C-terminal polypeptide fragment of the RBR1 protein that was expressed in E.coli as described earlier (Horváth et al., 2006). The purified recombinant protein was used to immunize rabbits by a company (Eurogentec). From the immunserum a crude IgG fraction was isolated by ammonium sulfate precipitation then IgG was further purified on protein gel blots of the antigen. To produce the PKL antibody, a C-terminal polypeptide encompassing the last 284 amino acids of the PKL protein has been expressed in fusion with a hexahistidine tag in the E.coli strain BL21DE3 Rosetta (Novagen). The protein was purified under denaturing conditions on Ni^2+^-nitrilotriacetic acid matrix (Ni-NTA Agarose, Qiagen) and used to raise polyclonal antibody in rabbits (Eurogentec). From the crude immunserum specific IgG fraction was isolated as described above.

### Microscopy

For BiFC assays Yellow Fluorescent Proteins (YFP) were visualized using a Leica TCS-SP2 laser scanning confocal microscope (LSM) (Leica Microsystems, Heidelberg, Germany) with a 63×1.4 NA oil-objective lens and processed using Leica confocal software v.2.61 about 16 h post-transformation. YFP was excited with a 514 nm argon laser and fluorescence detected between 520–550 nm. For *LBD16* promoter activity assays fluorescence detection of Green Fluorescent Proteins (GFP) was performed using a LSM 510, AxioObserver (Carl Zeiss, Germany) microscope with a 10×0.45 M27 objective lens. GFP was excited at 488 nm and the emitted light was captured at 505 to 555 nm. GFP fluorescence intensities captured by ZEN software (Carl Zeiss, Germany) were measured and quantified by ImageJ software (http://imagej.nih.gov/ij/).

### Transcript Level Analysis

Ten day-old seedling roots or suspension derived protoplasts were harvested, and total RNAs were extracted using the RNeasy kit (Qiagen). For quantitative RT-PCR, RNAs were treated with DNaseI (Ambion) and reverse transcribed using the first-strand cDNA synthesis kit (iScript Advanced cDNA Kit, BioRad). Gene-specific primers and iQSYBER Green Supermix (BioRad) were used on a C1000 Thermal Cycler (BioRad). Quantitative RT-PCR were performed using three replicates and At1g61670, At3g03210, At2g32760 were chosen as reference genes (Hruz et al., 2011; Vandesompele et al., 2002). Amplification cycles were analyzed using the Bio-Rad CFX Manager Software (Version 1.6) and fold expression change for each gene of interest was calculated using the ΔΔCt comparative method. Primer sequences are listed in Supplemental Table.

### Transient Expression of Proteins in Suspension Derived Protoplasts and Immunoprecipitation

For all experiments an *Arabidopsis* cell suspension derived from wild type *Col* roots (Mathur and Koncz, 1998) were used. Protoplast isolation and transient expression experiments were done according to (Fülöp et al., 2005), with slight modifications: 5×10^5^ protoplasts were used for each transformation with 3 to 5 μg plasmid DNA. After transfection, protoplasts were incubated in the dark for 16-24 h, harvested by centrifugation then proteins were extracted from the cell pellet as described previously (Cruz-Ramirez et al.,2012).

For the coimmunoprecipitation assays, 100 μg total protein extract was incubated in a total volume of 100 μl extraction buffer containing 150 mM NaCl and 1 μg anti-HA or 1.5 μg anti-cMyc or 2 μg anti-RBR1 antibody (Covance, clone 16B12 for HA and 9E10 for c-Myc, respectively). After 2 h, 10 μl Protein G-Sepharose matrix (GE Healthcare) was added which was previously equilibrated in TBS buffer and this mixture was further incubated for another 2 h on a rotating wheel at 4°C. The matrix was then washed in 3×500 μl washing buffer (1xTBS, 5% glycerol, 0,1% Igepal CA-630) and eluted by boiling in 25 μl 1.5x Laemmli sample buffer. Proteins were then resolved with SDS-PAGE and blotted to PVDF transfer membrane (Millipore). The presence of the proteins of interest was tested by immunodetection using rat anti-HA-peroxidase (3F10, Roche) or chicken anti-c-Myc primary antibody (A2128, Invitrogen), rabbit anti-chicken IgY HRP conjugate (Thermo Scientific) and anti-PKL antibody, respectively.

### Chromatin Immunoprecipiation (ChIP) Assays

ChIP analysis was performed as described (Gendrel et al., 2002; Nelson et al., 2006). Roots of 10 and 5 day-old *Arabidopsis thaliana* Columbia-0 and roots of 10 day-old *slr-1* and *ssl2-1* seedlings were harvested and proteins were cross-linked to DNA with 1% formaldehyde for 15 min. ChIP assays were performed using antibodies against PKL, RBR1 and normal rabbit IgG (Santa Cruz Biotechnology). Gene-specific primers and *SsoAdvanced SYBR Green* Supermix (BioRad) were used on a C1000 Thermal Cycler (BioRad). Quantitative ChIP PCR was performed in three replicates, and results were analyzed according to the percent input method or to the fold enrichment method (Haring et al., 2007). ChIP experiments were performed at least twice. Primer sequences are provided in Supplemental Table.

For plasmid ChIP, analysis protoplasts were prepared as described above and ChIP assays were performed as described in (Lavrrar and Farnham, 2004; Shaikhali et al., 2012).

## Acknowledgements

We thank Frederic Berger, Hidehiro Fukaki, Malcolm Bennett, Claudia Köhler, Jiri Friml for providing *pRBR1::RBR1-RFP*, *ssl2-1*, *slr-1, pPKL::PKL-GFP* seeds and the DR5 expressing vector, respectively. Authors are grateful to Hayashi Kenichiro for providing the auxinol compound and to Jürgen Kleine-Vehn and Rishi Bhalerao for stimulating discussions. The technical help of Adeline Rigal and Thomas Vain with the auxinol experiments is much appreciated. This work was supported by a postdoctoral fellowship of the Carl Tryggers Foundation (to KÖ) and by grants from Vetenskapsrådet and VINNOVA (to LB and SR).

