## Supplementary material for "PICKLE recruits RETINOBLASTOMA RELATED 1 to Control Lateral Root Formation in *Arabidopsis*"

### 1. Supplementary figures

**Supplementary Figure S1.** (A) Protein domain structure of the PKL protein showing the position of the LxCxE consensus pRb binding motives. (B) Protoplasts were transfected with *35S::HA-RBR1* (HA-RBR1) and *35S::myc-PKL-N* (Myc-PKL-N) and *35S::myc-PKL-C* (Myc-PKL-C) constructs and protein extracts from transformed protoplasts were immunoprecipitated with anti-myc antibodies. Immunocomplexes and unbound fractions were analyzed on protein gel blots using anti-HA or anti-myc antibodies, respectively. (C) Confocal microscopic images of the subcellular localization of the RBR1/PKL complex by BiFC assays. Coexpression of *35S::YFPN-RBR1* (RBR1) and *35S::YFPC-PKL-N* (PKL-N) or *35S::YFPC-PKL-C* (PKL-C) in suspension derived *Arabidopsis* protoplasts. PKL-N: N-terminal part of the PKL protein (aa 1-586), PKL-C: C-terminal part of the PKL protein (aa 463-1384), Bar = 5 μm. (D) BiFC assay in suspension derived protoplasts coexpressing RBR1 plus N-terminal part of PKL (PKL-N) versus RBR1 plus LxCxE mutated PKL-N (PKL-Nm) and RBR1 plus C-terminal part of PKL (PKL-C) versus RBR1 plus PKL-C LxCxE mutated (PKL-Cm) constructs.

**Supplementary Figure S2.** (A) Confocal images of the expression pattern of the *pRBR1::RBR1-RFP* (red channel) construct in different zones of the root (from right to left); root tip, the differentiation zone and during lateral root primordia formation. Bar represents 100 μm. White arrows point to pericycle-localized RBR1. (B) Confocal images of the expression pattern of the *pPKL::PKL-GFP* (green channel) construct in different zones of the root (from right to left); root tip, the differentiation zone and during lateral root primordia formation. Bar represents 100 μm. White arrows point to pericycle-localized RBR1 and PKL proteins.

**Supplementary Figure S3.** (A) 7 DAG wild type Columbia and *ssl2-1* seedlings. (B) Western blot analysis of RBR1 and PKL protein expression in wild type Columbia (Col), *ssl2-1* and *slr-1* roots probed with anti-RBR1 and anti-PKL antibodies, respectively.

**Supplementary Figure S4.** ChIP-PCR analysis of *LBD17*, *LBD18*, *LBD29* and *LBD33* promoter fragments. ChIP with the anti-RBR1 antibody using chromatin extracted from 10 DAG Col roots. Triangles show the position of the E2F consensus (E2F) and the AuxRE motives. Black bar represents the amplified PCR fragment position on the schematic diagram of the different promoters. -: non-template control, i: input DNA, m: IP with IgG, IP: IP with anti-RBR1 antibody.

**Supplementary Figure S5.** *LBD16* promoter activity (green channel) in wild type Columbia (Col-0), *amiRBR1* and *ssl2-1*. 6 DAG whole root confocal scans.

**Supplementary Figure S7.** 7 DAG wild type Columbia (Col-0), *amiRBR1*, *RBR1-RFP* and *ssl2-1* seedlings with and without Auxinol treatment.

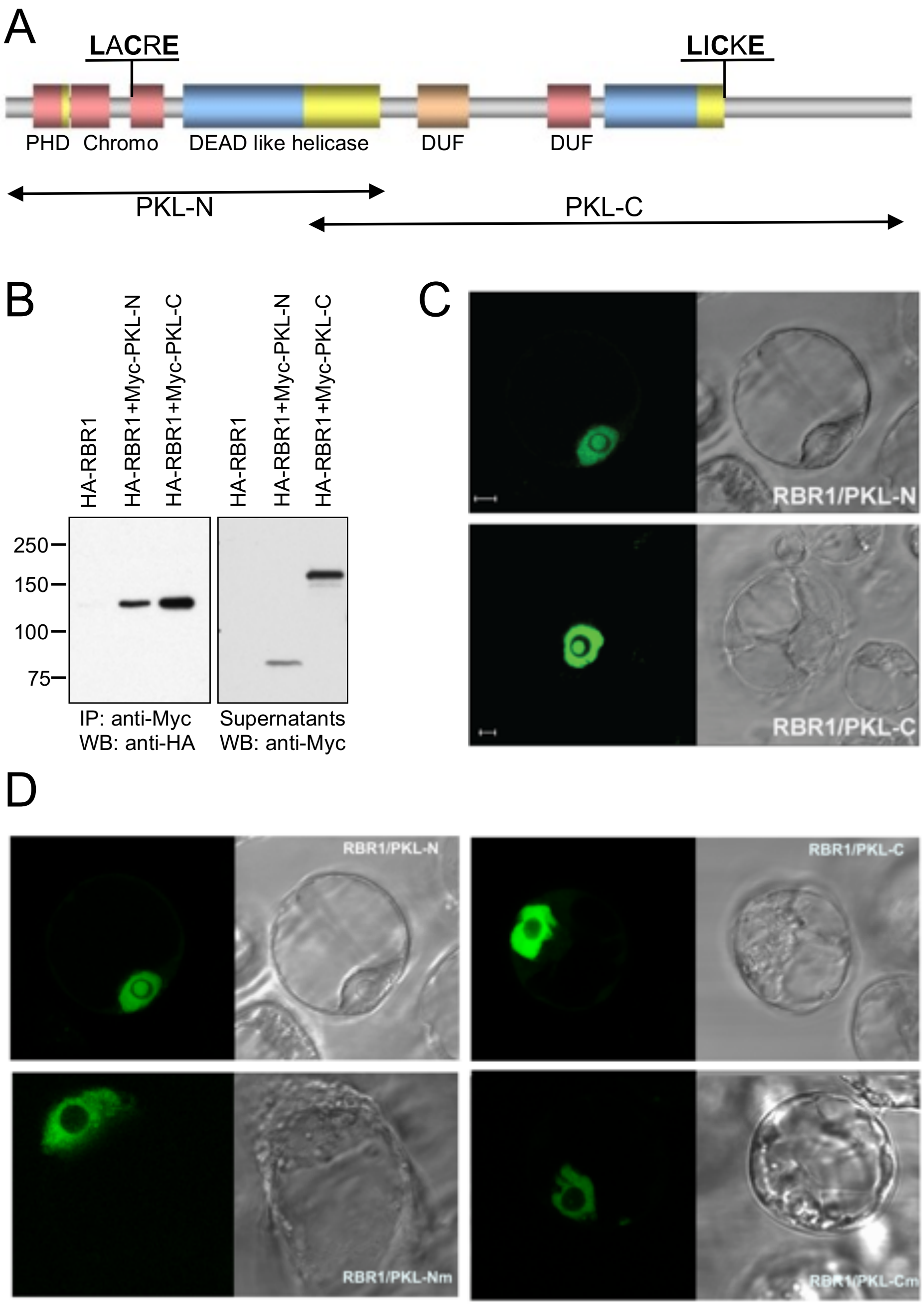


**Supplementary Figure S1.**

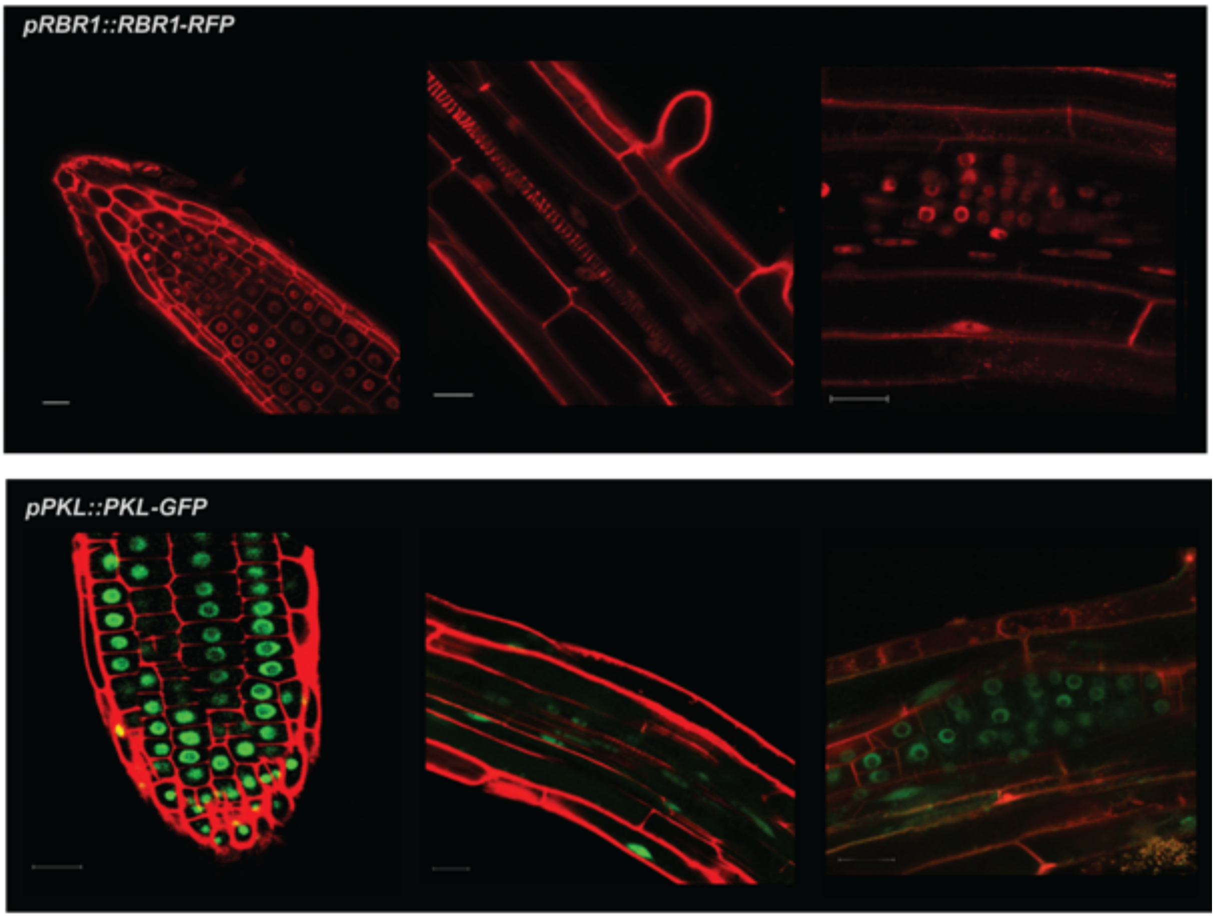


**Supplementary Figure S2.**

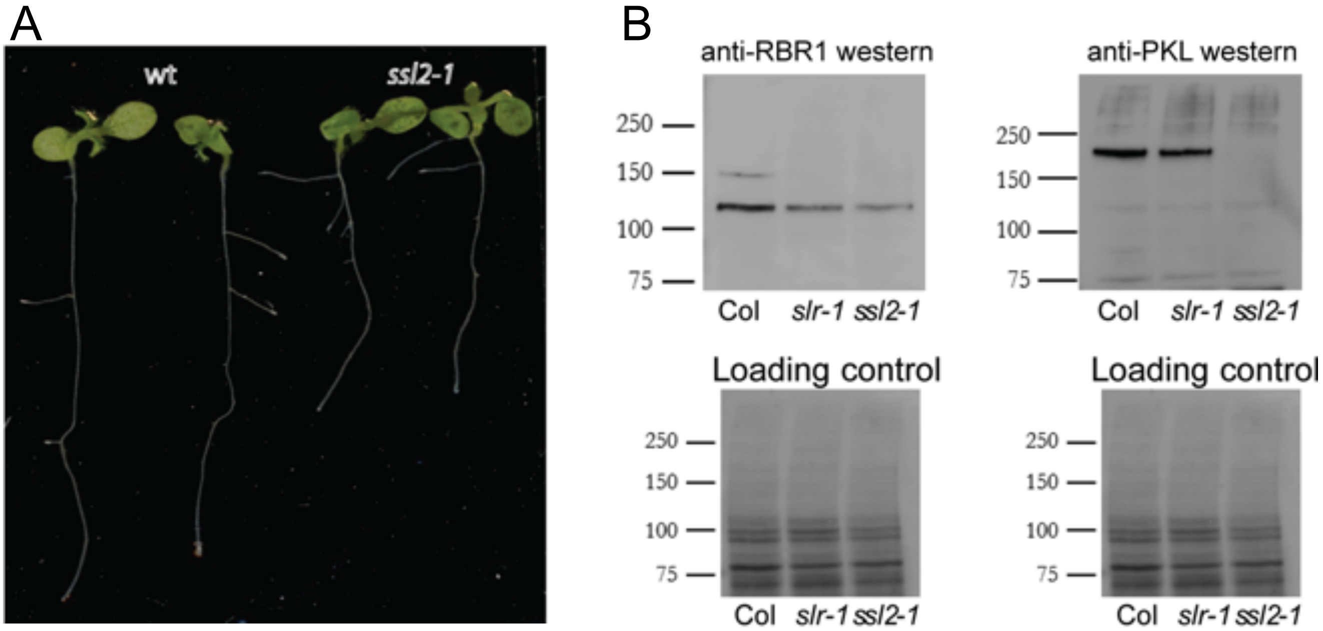


**Supplementary Figure S3.**

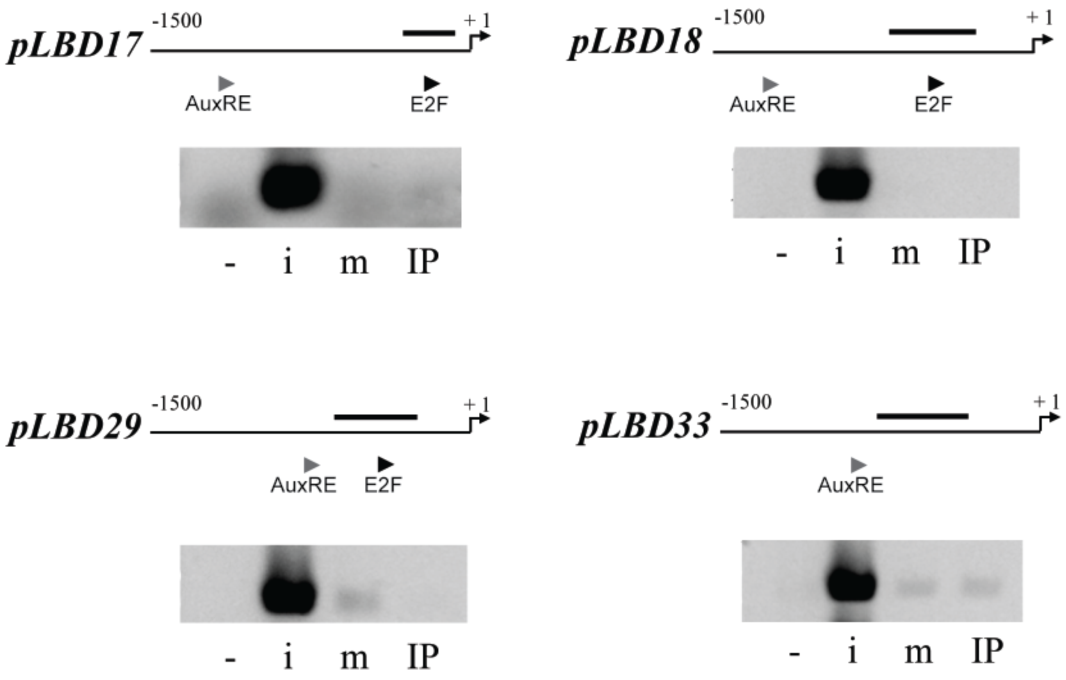


**Supplementary Figure S4.**

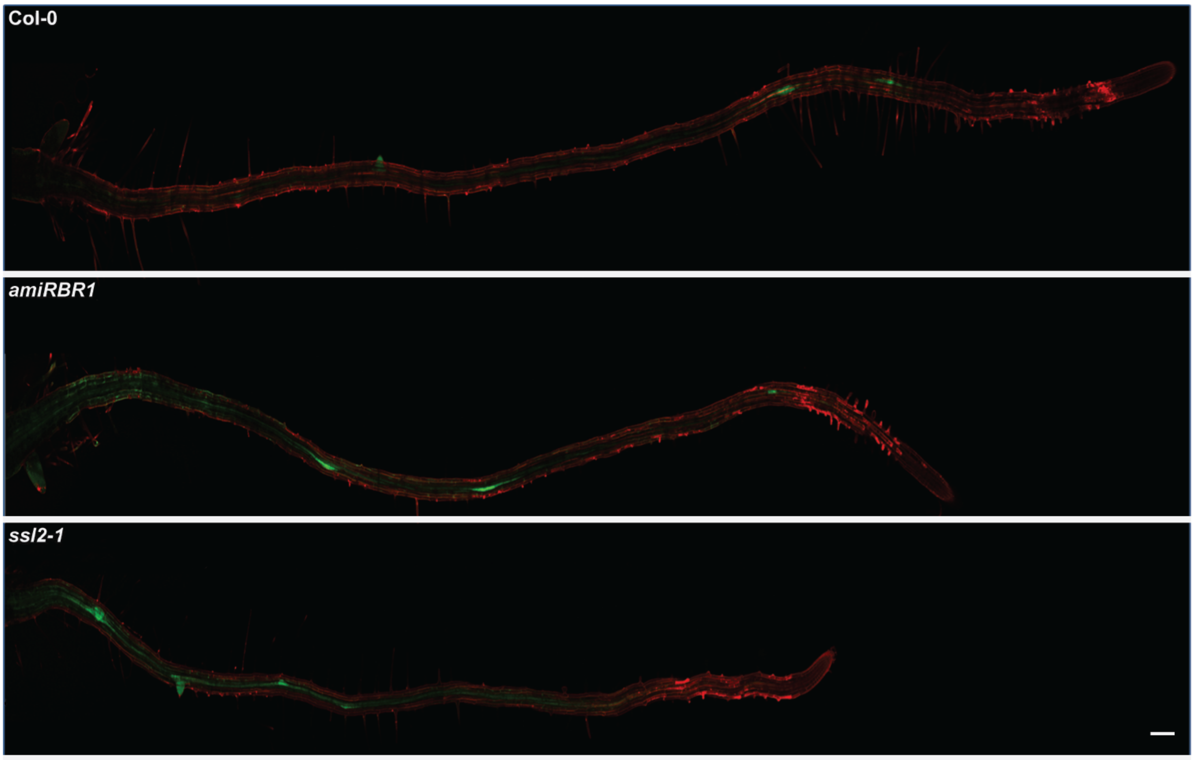


**Supplementary Figure S5.**

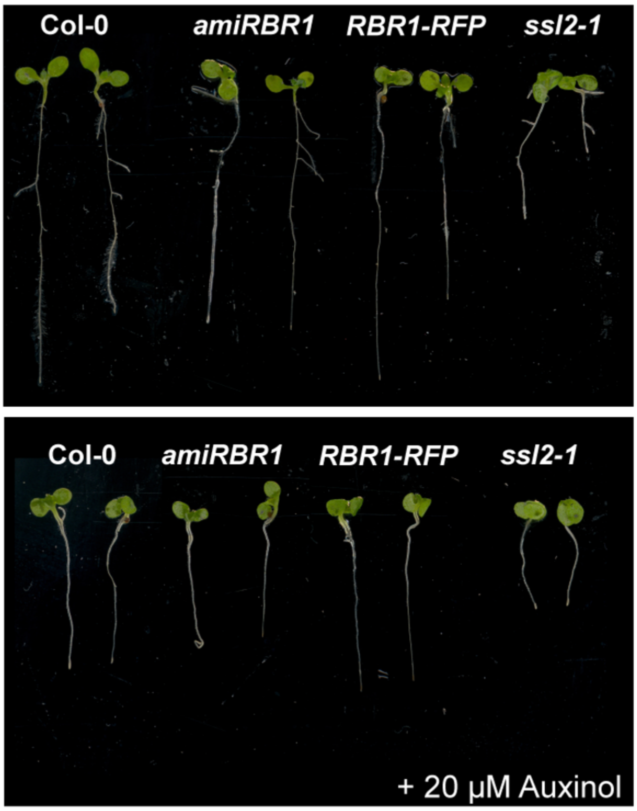


**Supplementary Figure S7.**

### 2. Supplementary Experimental Procedures

**Supplementary Table S1.**

List of oligonucleotide primers used in this study

The specificity of the PCR primers was assessed with the MFEprimer-2.0 program [1,2].

| | |
|---|---|
| **qPCR primers** | |
| RBR1 | GGATCGCCAAAGGTGTCTGTGTTTC<br>TCCTCTTGCGGTTGCTGCTGCTGTT |
| LBD16 | GCCACAGAGCTCGCTAGAAGA<br>TCGCACGTTGGCTGTTGT |
| ACR4 [3] | GATCATAGTGCGGTCTGTTGG<br>AGGGATAGAAGCAGGGAAACC |
| At1g61670 (Ref01) | CTGACCCTCTAATCGAACTC<br>CCCTGTTTCCGCTAAGTATG |
| At3g03210 (Ref04) | CATGGGAACTGGGTTATGG<br>CTCCTTCTTGCTCCTTTGG |
| At2g32760 (Ref02) | CCGACAAGAAAGAAGTTCAG<br>GATAACGAATTGGCACTCTC |
| **ChIP primers** | |
| LBD16 F1 | CCACACGTGCCTAAGCCACCTAAGCAGA<br>GCACTTTGCAAGTTTGAAACTC |
| LBD16 F2 | GTGGCTTCTCCTCAAATGTC<br>ACGAGCGAACTAGGAAATGG |
| LBD16F3 | GCTCCAAGAACCAAACGTATCC<br>GCACTTTGCAAGTTTGAAACTC |
| LBD16 F4 | TCGCTCGTGCACATCTCCGA<br>TGTCCCCGCCGTTGTACCGT |
| LBD17 | GAAAGGTGTAGGCCACTAAG<br>GAATGGAGGAGCATTAGC |
| LBD18 | GATAAGTTGGCGAACGAGTC<br>GCGTGAAGCAAACATGCGATAAGC |
| LBD29 | TCTCTGGAAGGGTAGGTTAG<br>GGTGAAGATGAAGGTGGGTTAC |
| LBD33 | GTGTGGATCGGAGTAATG<br>GCATCGTGGCCTACATTT |
| ***pLBD16::GFP* plasmid ChIP primers** | |
| genomic specific | TCGCTCGTGCACATCTCCGA<br>TGTCCCCGCCGTTGTACCGT |
| plasmid specific | CACCCAAAGACACCAAACAC<br>AAGATGGTGCGCTCCTGGAC |
| **Site directed mutagenesis primers** | |
| PKL-N SDM BspFwd | AGATCTCGAGCTCAAGCTTCGAATTCTG |

| | |
|---|---|
| PKL-N SDM Fwd | ATCGGATTGCTGCCGCCCGGGCGGAAG |
| PKL-N SDM PspXRev | ATGTTCTCTGATAACTGCTCGAGCTTG |
| PKL-N SDM Rev | CTTCCGCCCGGGCGGCAGCAATCCGAT |
| PKL-C SDM SpeIFwd | GGAAGATGTGAGCTCTGATGG |
| PKL-C SDM Fwd | GACAAAGAGTTGGGGATCCAAGAGGCTATCGCCAAGGCCTTGAATTTCCCTCACATAAGTTTG |
| PKL-C SDM BclIRev | GTAATACGACTCACTATAGGGCG |
| PKL-C SDM Rev | CAAACTTATGTGAGGGAAATTCAAGGCCTTGGCGATAGCCTCTTGGATCCCCAACTCTTTGTC |

**Generation of plasmid constructs**

For co-immunoprecipitation (Co-IP) assays, the epitope tagging vectors containing 3x*Myc* or 3x*HA* tags, *pRT104-3Myc* and *pRT104-3HA*[4] vectors and for *Bimolecular Fluorescence Complementation (BiFC) assays pSAT1-cEYFP-C1-B* (*35S::YFPC*) and *pSAT1-nEYFP-C1* (*35S::YFPN*)[5] vectors were used. All constructs contain the epitopes at the N terminus of the fusion proteins. The coding sequence of RBR1 was PCR amplified from cDNA using RBR1for/*BamH*I (ATAGGATCCATGGAAGAAGTTCAGCCT) and RBR1rev/*Xho*I (ATACTCGAGCTATGAATCTGTTGGCTC) primer pair. The *BamH*I and *Xho*I digested PCR product was inserted into the *Sal*I and *BamH*I digested *pRT104-3HA* vector to create HA-RBR1. Myc-PKL-C and Myc-PKL-N contains the C-terminal part (aa 463-1384) and the N-terminal part (aa 1-586) of the PKL protein as *KpnI* and blunt ended *Hind*III inserts respectively. The *35S::YFPN-RBR1* construct was obtained by excising the coding region of RBR1 from the HA-RBR1 construct using *BamH*I and *Apa*I and inserting it into the *Apa*I and *Bgl*II sites of the *pSAT1-nEYFP-C1* vector. The coding region of the full length PKL (fPKL) protein was assembled by inserting the synthetized *Sac*II/*Hind*III digested N-terminal part into the *Hind*III/*Sac*II sites of the modified *pBluecsript* vector containing the C-terminal part of the protein (*pmBluescript2-PKL-C*) provided by Riken to create *pmBluescript2-fPKL*. The *35S::YFPC-fPKL* construct contains fPKL as an *Sac*II/*Kpn*I blunt ended insert. The *35S::YFPC-PKL-N* construct was generated by ligating the *Hind*III fragment of the *pmBluescript2-fPKL* construct into the *Hind*III site of the *pSAT1-cEYFP-C1-B* vector. The *35S::YFPC-PKL-C* was obtained by ligating the blunt ended *Sfi*I PKL-C insert from *pmBluescript2-PKL-C* into the blunt ended *Kpn*I site of the *pSAT1-cEYFP-C1-B* vector.

Mutation of the LxCxE motives into AxAxA in the N- and C-terminal part of the PKL protein was performed by PCR-based site directed mutagenesis. For primer sequences used for the mutagenesis see Supplemental Table.

For promoter activity assays, constructs in the binary vector *pSL36-GFP* were used. The *pLBD16::GFP* construct was generated by PCR amplification of a 1551 bp promoter fragment upstream of the translational initiation site using primers LBD16for/*Pml*I (CCACA CGTGCCTAAGCCACCTAAGCAGA) and LBD16rev/*Spe*I (TGGACTAGTGCGAAACGAACAAAAAAG). The PCR fragment was digested using *Pml*I and *Spe*I and inserted into the *Pml*I/*Spe*I digested *pSL36-GFP* vector. Similarly the *pLBD29::GFP* was obtained by PCR amplification of a 1456 bp promoter fragment upstream of the translational initiation site using primers LBD29for/*Pml*I (CCACACGTGGTCCATTGTCTCTATATATCG) and LBD29rev/*Xba*I. The PCR fragment was digested with *Pml*I/*Xba*I and ligated into the *Pml*I/*Spe*I digested *pSL36-GFP* vector. *pLBD16Δ1::GFP* and *pLBD16Δ2::GFP* are deletion variants of the original *pLBD16::GFP* construct, missing a 738 bp *Pml*I - *Bst*1107I and a 813 bp *Bst*1107I - *Spe*I promoter fragment, respectively.

For *DR5* activity assays, the *DR5* sequence was amplified by PCR with DR5fwdPmlI (TACACGTGAAATAGGCGTATCACGAGGC) and DR5revSpeI (ATACTAGTGATCGATCCCCTGTAATTGT) primer pair from the *pDR5rev::GFP* construct[6]. The PCR fragment was digested with *Pml*I/*Spe*I and ligated into the *Pml*I/*Spe*I digested *pSL36-GFP* vector.
